# Spatial density estimates of Eurasian lynx (*Lynx lynx*) in the French Jura and Vosges Mountains

**DOI:** 10.1101/600015

**Authors:** Olivier Gimenez, Sylvain Gatti, Christophe Duchamp, Estelle Germain, Alain Laurent, Fridolin Zimmermann, Eric Marboutin

## Abstract

Obtaining estimates of animal population density is a key step in providing sound conservation and management strategies for wildlife. For many large carnivores however, estimating density is difficult because these species are elusive and wide-ranging. Here, we focus on providing the first density estimates of the Eurasian lynx (*Lynx lynx*) in the French Jura and Vosges mountains. We sampled a total of 413 camera trapping sites (with 2 cameras per site) between January 2011 and April 2016 in seven study areas across seven counties of the French Jura and Vosges mountains. We obtained 592 lynx detections over 19,035 trap days in the Jura mountain and 0 detection over 6,804 trap days in the Vosges mountain. Based on coat patterns, we identified a total number of 92 unique individuals from photographs, including 16 females, 13 males and 63 individuals of unknown sex. Using spatial capture-recapture (SCR) models, we estimated abundance in the study areas between 5 (SE = 0.1) and 29 (0.2) lynx and density between 0.24 (SE = 0.02) and 0.91 (SE = 0.03) lynx per 100 km^2^. We also provide a comparison with non-spatial density estimates and discuss the expected discrepancies. Our study is yet another example of the advantage of combining SCR methods and non-invasive sampling techniques to estimate density for elusive and wide-ranging species, like large carnivores. While the estimated densities in the French Jura mountain are comparable to other lynx populations in Europe, the fact that we detected no lynx in the Vosges mountain is alarming. Connectivity should be encouraged between the French Jura mountain, the Vosges mountain and the Palatinate Forest in Germany where a reintroduction program is currently ongoing. Our density estimates will help in setting a baseline conservation status for the lynx population in France.

## Introduction

Obtaining estimates of animal population density is a key step in providing sound conservation and management strategies for wildlife [1]. For many large carnivores however, estimating density is difficult because these species are elusive and wide-ranging, resulting in low detection rates [2]. To deal with these issues, non-invasive techniques, such as camera trapping and DNA sampling, are increasingly used [3]. These non-invasive techniques generate data that can be analyzed with capture-recapture methods to estimate densities [4].

Standard capture-recapture models for closed populations [5] have long been used to estimate animal abundance and density, including many large carnivores [6,7]. However, when converting abundance into density, density estimates are highly sensitive to the size of user-defined area assumed to reflect the effective sampling area [8]. In addition, individual heterogeneity on the detection due to spatial variation in the distance of home ranges to the sampling devices may lead to biased density estimates [5]. Spatial capture-recapture (SCR) models deal with these issues by explicitly incorporating spatial locations of detections [9–12], and they are increasingly used to estimate densities of large carnivores [13–18].

Here, we focus on the threatened Eurasian lynx (*Lynx lynx*) in the French Jura and Vosges mountains (see [19] for a map of its distribution in Europe; see also https://www.lcie.org/Large-carnivores/Eurasian-lynx for recent updates). As in many regions of western Europe [20], lynx were extirpated from France between the 17^th^ and 20^th^ centuries due to habitat degradation, persecution by humans and decrease in prey availability [21]. Shortly after their initial reintroduction in Switzerland in the 1970s [22], lynx naturally increased their range and started recolonizing France by repopulating forests on the French side of the Jura [21]. Reintroductions also occurred in the French Vosges mountain between 1983 and 1993 with the perspective of establishing a population there [23]. The species is listed as endangered in the IUCN Red list and is of conservation concern in France due to habitat fragmentation, poaching and collisions with cars and trains. Currently, the French population of lynx is restricted to three mountain ranges, the Vosges, in northeastern France, the Jura and the Alps, with little connectivity between them most likely due to human-made linear infrastructures. While the Northern Alps are slowly being recolonized with lynx mostly coming from the Jura [24], the Jura holds the bulk of the French lynx population. In contrast, the lynx presence in the Vosges mountain remained stable following the reintroductions and then, has been continuously decreasing since 2005 [25].

Despite their conservation status, little information on abundance and density of lynx in France exist. In this study, we used SCR and standard capture-recapture models to provide the first estimate of lynx abundance and density using camera-trap surveys implemented in the French Jura and Vosges mountains from 2011 to 2016. Based on these results, we discuss research and management priorities for the effective conservation of lynx in France.

## Methods

### Ethics statement

We used non-invasive methods for data collection, which did not involve manipulation or handling of any living organism. Therefore, approval from an animal ethics committee was not required. Cameras were set on public or private forests with the permission of local authorities or local owners, respectively. We advertised the study and the presence of camera traps to the local stakeholders and the public visiting the areas. In agreement with French legislation, we deleted photos permitting the identification of goods or people.

### Study area and sampling design

The study area encompassed three counties of the French Jura mountain, namely Ain, Doubs and Jura and four counties of the Vosges mountain, namely Vosges, Haut-Rhin, Bas-Rhin and Moselle (Figure 1). Elevation ranged from 163 and 1,718 m above sea level in the Jura mountain and from 104 to 1,422 m in the Vosges mountain. The human population density was 88 per km^2^ in the Jura mountain and 170 per km^2^ in the Vosges mountain. Forests cover 50% on average of the Jura mountain [26] and 70% of the Vosges mountain [27]. Sampling occurred over 6 years, between January 2011 and April 2016, mostly in winter and spring, with surveys lasting between 2 and 4 months. We considered two study areas in 2011, 2014 and 2015, three study areas in 2013 and one study area in 2012 and 2016 through camera trapping (Figure 1).

**Figure 1:**
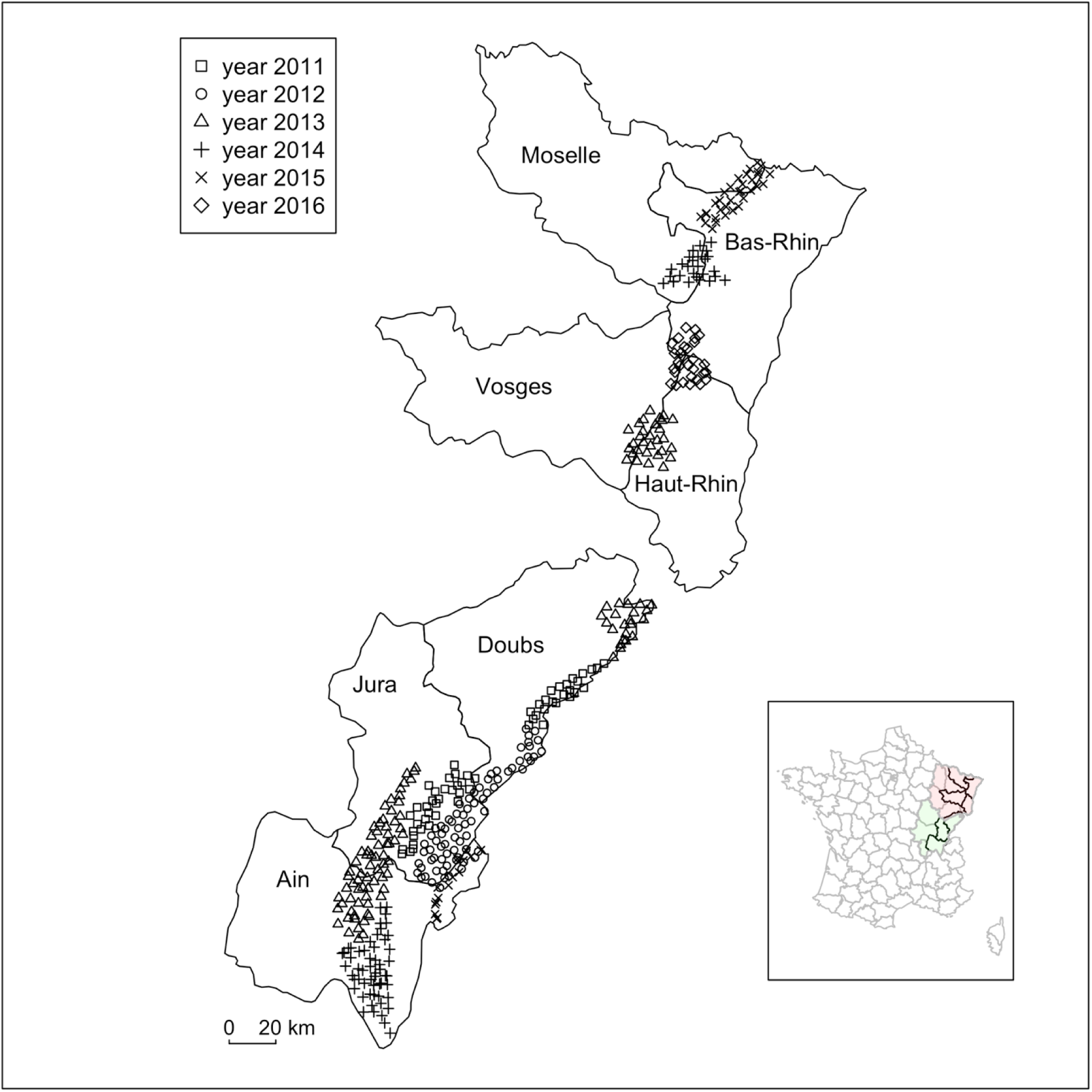
Map of the study area in the French Jura and Vosges mountains. The study area encompassed seven counties (Ain, Jura and Doubs in the Jura mountain and Vosges, Haut-Rhin, Bas-Rhin and Moselle in the Vosges mountain) that were monitored through 418 camera trapping sites (298 in the Jura mountain and 115 in the Vosges mountain; two camera traps were set per site), each within a 2.7 × 2.7 km cell. The inset map represents the French counties (grey borders), the counties that were considered in the study (black borders), the Jura mountain (green shaded area) and the Vosges mountain (red shaded area).

We divided each study area into a grid of 2.7 × 2.7 km cells applying a systematic design where one out of two cells was sampled [28], hence ensuring that at least one camera trap was set in each potential lynx home range (between 100km^2^ and 250km^2^, see [29]). To maximize detectability, we set (non-baited) camera traps in forested habitats, based on previous signs of lynx presence and on local knowledge, at optimal locations where landscape and terrain features were likely to channel lynx movements on more predictable paths (on forest roads, hiking trails and to a lesser extent on game paths) [30]. Camera were settled on within a single session design continuously during 60 days between February and beginning of March with little variation between sites.

At each trapping location, we set two Xenon white flash camera traps (models: Capture, Ambush and Attack; Cuddeback, WI, USA) with passive infrared trigger mechanisms to photograph both flanks of an animal. We checked camera traps weekly to change memory cards, batteries and to remove fresh snow after heavy snowfall. Based on unique coat patterns, we identified individual lynx on photographs [31]. The recognition of individual was computer-induced, not fully automated. We used the Extract-compare © software that compares the lynx spot pattern with a library of previously extracted pattern and proposes potential matches according to a score (http://conservationresearch.org.uk/Home/ExtractCompare). The observer can confirm the lynx identification or not and browse through the highest-ranking proposed matches. The final decision is made by the observer based on an additional visual examination of the entire photo set for this particular lynx. Pictures for which no match was found with the software were visually checked against our entire photo library. Only when the match is undeniable is the individual recorded as a match, otherwise it was recorded as a new individual. All captures that do not fit automated or associated visual confirmation with no doubt, because of a poor picture quality (e.g. blurry, overexposed), were classified as “unconfirmed” and excluded from the analyses. We recorded the date, time, sex whenever possible, and location of each photographic capture of a lynx. During the time of year our study took place, juvenile lynx (< 1 year old) can still be with their mother [32]. In our analysis, we retained only independent lynx, i.e. adult lynx or emancipated individuals based on physical characteristics or previous knowledge of their age or status (from photographic evidence). We defined a capture occasion as 5 successive trap nights [30], dissociating trapping events from individual photo to avoid pseudo-replications.

### Spatial capture-recapture analyses

We used spatial capture-recapture (SCR) models to estimate lynx densities [4]. In contrast with standard (non-spatial) capture-recapture models, SCR models use the spatial locations of captures to infer the activity center (or home range) of each individual. We assumed that individual encounters are Bernoulli random variables with individual- and trap-specific detection probabilities. More precisely, the detection probability *p*_*ij*_ of an individual *i* at trap *j* is assumed to decrease as the distance (*d*_*ij*_) from its activity center increases according to a detection function. We used the half-normal detection function, *p*_*ij*_ = *p*_*0*_ exp(-d_ij_^2^/(2σ^2^)), where *p*_*0*_ is the probability of detecting an individual when the trap is located exactly at its center of activity and σ is the spatial scale (or movement) parameter that controls the shape of the detection function. For one of the two study areas in the French Jura mountain in years 2011 and 2013, we detected only a few individuals (see the columns Doubs in Table 1). To increase the effective sample size, we combined the data from the two sampling areas using common detection and spatial parameters for both areas, while estimating density separately (e.g., [33]). We defined a state-space, i.e. the area encompassing all potential activity centers of the observed individuals, by building a grid that buffered outermost camera trap locations by 15 km (corresponding to at least 2σ [4]) with a resolution of 1.5 km (or pixels of area 2.25 km^2^). We fitted SCR models in the maximum likelihood framework using the R package oSCR [34,35].

**Table 1:** Main characteristics and results of the lynx camera-trap survey carried out in a) the French Jura mountain and b) the French Vosges mountain.

| **a. Year/County** | 2011/Doubs | 2011/Jura | 2012/Jura & Doubs | 2013/Doubs | 2013/Ain & Jura | 2014/Ain | 2015/Ain |
| --- | --- | --- | --- | --- | --- | --- | --- |
| **Period of trap activity** | Jan-Apr | Feb-Apr | Feb-Apr | Feb-Apr | Feb-Apr | Feb-Apr | Feb-May |
| **Number of active camera traps** | 48 | 66 | 148 | 44 | 142 | 118 | 30 |
| **Number of trapping days (average/area)** | 63 | 59 | 69 | 63 | 58 | 59 | 99 |
| **Number of capture occasions<sup>i</sup>** | 15 | 15 | 17 | 14 | 13 | 13 | 21 |
| **Number of detections** | 22 | 42 | 130 | 25 | 117 | 158 | 38 |
| **Number of detected individuals** | 4 | 9 | 21 | 6 | 19 | 23 | 10 |
| **Number of females, unknown, males** | 1, 1, 2 | 1, 7, 1 | 2, 14, 5 | 1, 4, 1 | 2, 13, 4 | 4, 16, 3 | 2, 8, 0 |
| **Number of detections / ind: mean, min, max** | 3, 2, 4 | 2.8, 1, 6 | 2.5, 1, 10 | 2.7, 1, 6 | 3.6, 1, 11 | 3.3, 1, 9 | 2.2, 1, 5 |

| **b. Year/County** | 2013/Haut-Rhin & Vosges | 2014/Bas-Rhin & Moselle | 2015/Bas-Rhin & Moselle | 2016/Bas & Haut-Rhin |
| --- | --- | --- | --- | --- |
| **Period of trap activity** | Dec-Jan | Feb-Apr | Feb-Apr | Feb-Apr |
| **Number of active traps** | 60 | 50 | 60 | 60 |
| **Number of trapping days (average/area)** | 52 | 59 | 57 | 59 |
| **Number of detections** | 0 | 0 | 0 | 0 |

<sup>i</sup>A capture occasion is defined as 5 successive trap days.

For comparison, we also estimated abundance using standard (non-spatial) capture-recapture models [5]. We dropped the spatial information and considered only the detections and non-detections for each individual. We considered two models, M0 in which the detection probability is the same for all individuals, and Mh in which the detection probability varies among individuals. We fitted standard models in the maximum likelihood framework using the R package Rcapture [36]. We estimated density as the ratio of estimated abundance over an effective trapping area (ETA). ETA was estimated by adding a buffer to the trapping area equal to the mean maximum distance moved (MMDM) or half of it (HMMDM). We calculated the MMDM by averaging the maximum distances between capture locations for all individuals detected at more than one site.

## Results

We collected data from 413 camera trapping sites (2 camera traps were set per site) resulting in 25,839 trap days (Table 1). In total, we identified 92 lynx over 532 detection events in the Jura mountain, including 16 females, 13 males and 63 individuals of unknown sex. The number of detections per individual was 2.6 on average and varied from 1 up to 11. In contrast, we collected no lynx photo in the Vosges mountain, therefore we did not proceed with analyses for this area.

For the Jura mountain, abundance estimates were similar whether we used spatial or non-spatial models, although always slightly higher for the former. Estimated abundance among study areas varied between 5 (SE = 0.1) and 29 (0.2) lynx in the spatial analyses, between 4 (0.7) and 23 (0.7) with model M0, and between 5 (1.7) and 28 (3.6) with model Mh. Estimated density varied between 0.24 (0.02) and 0.91 (0.03) lynx per 100 km^2^ in the spatial analyses (Table 2). In the non-spatial analyses, the density varied between 0.31 (0.05) and 0.78 (0.02) lynx per 100 km^2^ under model M0 and between 0.34 (0.06) and 0.95 (0.12) under model Mh when the MMDM was used. When we used HMMDM, the density varied between 0.57 (0.10) and 1.46 (0.16) lynx per 100 km^2^ under model M0 and between 0.67 (0.12) and 1.43 (0.16) under model Mh.

**Table 2:**
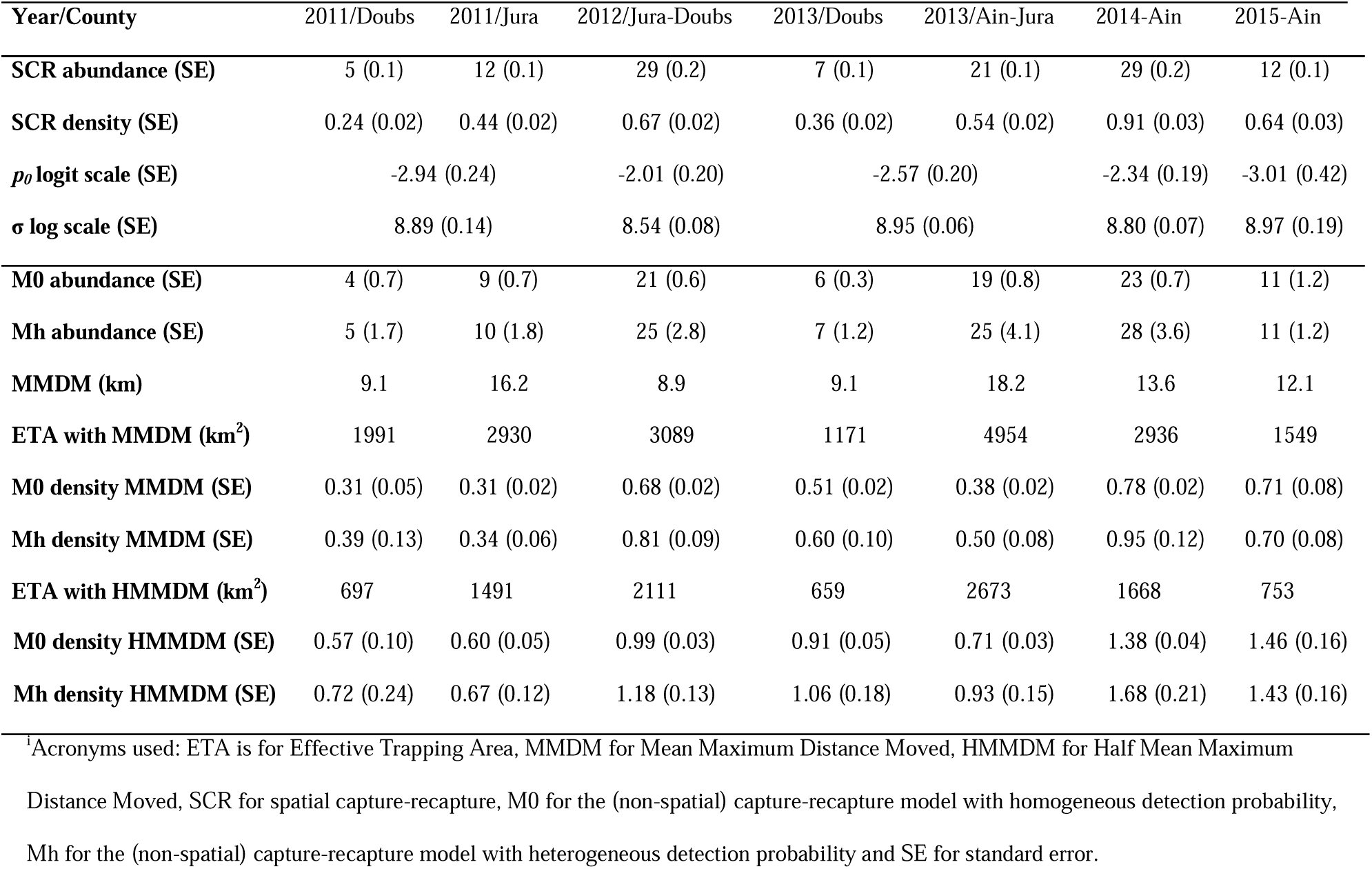
Lynx abundance and density estimates obtained from spatial and non-spatial capture-recapture analyses of camera-trapping data collected in the French Jura mountain. Densities are provided in number of lynx per 100 km^2^. For 2011 and 2013, parameters of the spatial capture-recapture model (*p*_*0*_ and σ) are common to both areas in each year. Acronyms are defined in the footnote^i^.

From the spatial analyses, we used the model estimates to produce density surfaces within the state-space (Figure 2). The density per pixel of area 2.25 km^2^ ranged from 0 to 0.20 individuals in the Jura mountain.

**Figure 2:**
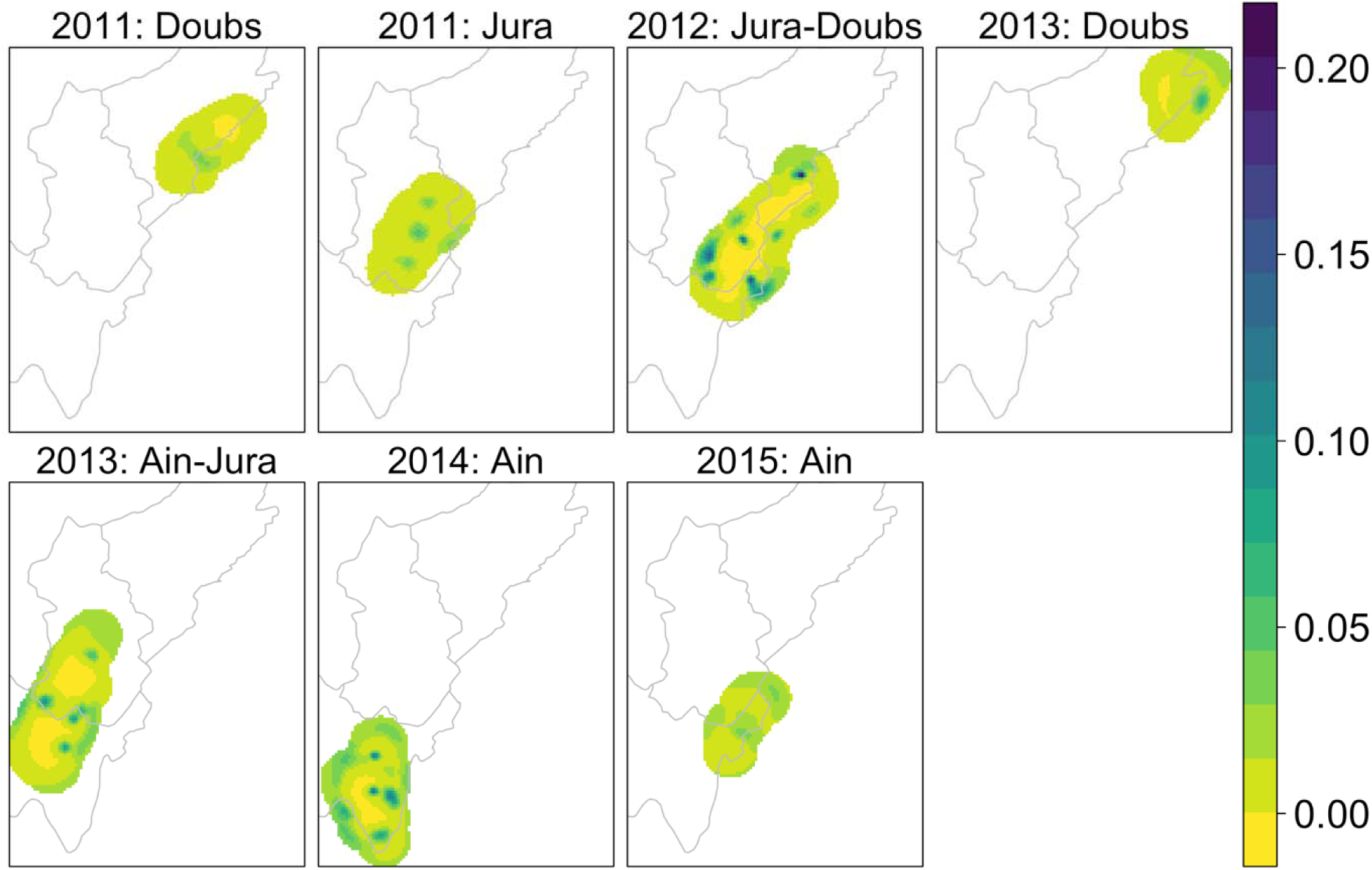
Lynx (*Lynx lynx*) density maps in the French Jura mountain. The density scale is in lynx per 2.25 km^2^ (pixel resolution is 1500m × 1500m). We obtained the estimated abundance in each map by summing up the densities in each pixel altogether. Yellow is for low densities, green for medium densities and blue for high densities; the density scales are specific to each map. Note that the interpretation of these plots as density maps is subject to caution (see the vignette “secr-densitysurface” of the SECR R package [58]).

## Discussion

By using camera-trap sampling and SCR models, we provided the first multi-site density estimates for lynx that will help in setting a baseline conservation status for the French lynx population. The multi-site dimension of our study allows exploring variability in the density estimates across landscapes. Our study is yet another example of the potential of combining SCR methods and non-invasive sampling techniques to estimate abundance and density for elusive and wide-ranging species, like large carnivores [13–18].

When examining densities across study areas in the French Jura mountain, we found spatial variation between the three counties, with Doubs area having the lowest densities, Ain the highest densities, and Jura intermediate densities. Our density estimates were of similar magnitude to other lynx populations in Europe: 1.47 and 1.38 lynx / 100 km^2^ in the Northwestern Swiss Alps [13], 0.58 (Štiavnica mountains) and 0.81 individuals / 100 km^2^ (Velká Fatra National Park) in Slovakia [37] and 0.9 individuals / 100 km^2^ in the Bavarian Forest National Park in Germany [38].

While [13] and [37] used SCR models, [38] used standard capture-recapture models with HMMDM to estimate densities, which makes them difficult to compare [39]. Indeed, in other carnivore studies, the use of HMMDM also produced similar density estimates to SCR models [13], while in others, including ours, the SCR estimates were closer to the MMDM estimates [2] or intermediate between the MMDM and HMMDM estimates [40]. When looking at reference values for densities across the distribution range of the species, it may be biologically meaningful to use the MMDM density estimate as a reference as it covers the whole potential of animal movements. On the other hand, because SCR models make space explicit whereas standard model-based densities are sensitive to the definition of the effective sampling area, we recommend the use of SCR models to estimate lynx densities. Our lynx density estimates might suffer from potential sources of bias that need to be discussed. First, the period of sampling is important to account for when setting up camera trap surveys [41]. We conducted our survey outside the dispersal period, during the lynx mating season (February-March mostly). We did so to avoid capturing transient individuals and to increase detectability because of high lynx activity and relatively reduced human activities [31].

However, some individuals might have moved in and out of the study areas, especially males who cover greater distances during the mating season. Whereas the presence of non-resident individuals can affect the calculation of (H)MMDM, and in turn density estimated with standard capture-recapture models, SCR density estimates were found to be robust to the presence of transient individuals [42]. Second, males have larger home ranges than females [13], which leads to heterogeneity in the SCR model parameter estimates. Because there were too few males and females identified and lots of individuals with unknown sex, sex-specific SCR analyses [43] produced unreliable abundance and density estimates (results not shown). If detection heterogeneity is ignored in capture-recapture models, abundance is underestimated [44], therefore our density estimates are probably biased low and should be considered as a conservative metric. The determination of sex could be improved by i) combining the photographic surveys with genetic surveys, ii) conducting deterministic surveys over several years (e.g., [13]), iii) conducting an opportunistic camera trapping survey all over the years and setting camera trap at fresh lynx kills, iv) setting infrared flash camera traps capable of taking burst of images in rapid sequence at marking sites regularly used by the lynx (e.g., [45]). Last, we did not detect any individuals in the Vosges mountain, even though the sampling effort was similar to that implemented in the Jura mountain (Table 1). This finding is likely to be representative of the current critical situation of the lynx in the Vosges mountain.

We envision several perspectives to our work. First, while density estimates are of primary interest for conservation, understanding the mechanisms underlying trends in abundance is required to make sound conservation decisions [1]. SCR models have been extended to open populations [46] and can be used to estimate demographic parameters (survival, reproduction) of large carnivores [47]. Unfortunately, because of logistic constraints, we could not sample the same areas over several years, which precludes a standard application of these models. A solution may lie in the combination of the data we collected through systematic camera-trap surveys with additional data in the SCR framework, such as occupancy data [48] or opportunistic camera-trap data [49]. Second, in addition to traffic-induced mortality and conflicts with human activities, the expansion of lynx populations is limited by habitat fragmentation [50], hence the need to assess connectivity with other populations [51]. SCR models can be used to quantify landscape connectivity by replacing the Euclidean distance between camera traps and home range centers by the least-cost path [52,53]. For lynx, this will require setting up traps across a gradient of habitat types, not only forested habitats, so that resistance to movement can be estimated.

In conclusion, our lynx density estimates for the French Jura mountain complement nicely the estimates recently provided for the Northwestern Swiss Alps [13]. The use of camera-trapping coupled with SCR models in both France and Switzerland was the result of a cooperation between the two countries with the perspective of a transboundary monitoring [54,55]. This approach would prove useful to accurately estimate densities in other areas where habitats and prey availability might differ, and overall lynx detectability varies. Also, collecting and adding movement data from GPS-collared lynx would be useful [49,56] to try and infer the connections between subpopulations.

The case can be made for monitoring the return of the lynx in the French Alps. Indeed, small-scale camera-trapping surveys and opportunistic observations are currently active and producing signs of lynx presence. However, the lack of a coordinated and intensive sampling effort prevents us from being able to estimate abundance and density and inferring trends.

In contrast, the situation in the Vosges mountain is alarming with no individuals detected over the study period. Because the Vosges mountain are located between the French Jura mountain and the Palatinate Forest in Germany where a reintroduction program is ongoing (program LIFE13 NAT/DE/000755), the lynx colonization in the Vosges mountain remains possible both by the north and the south. Incidentally, two cases of lynx dispersal in the Vosges mountain from neighboring mountains have been recently observed ([57]; program LIFE13 NAT/DE/000755). To ensure the detection of lynx in the Vosges mountain, we recommend reinforcing collaborative monitoring by involving all field stakeholders and enhancing communication on the species signs of presence.

In this context, obtaining accurate and comparable lynx densities will be crucial to closely monitor population trends at the national scale and inform management policies for the effective conservation of the Eurasian lynx in France.

## Acknowledgments

We thank the staff from the French National Game and Wildlife Agency (ONCFS), the CROC, the Forest National Agency, the “Directions Départementales des Territoires”, the “Fédérations Départementales des Chasseurs” the Regional Natural Parks, the environmental protection associations and all the volunteers from the “Réseau Loup Lynx” who collected the photographs during the camera-trapping session. OG was funded by CNRS and the “Mission pour l’Interdisciplinarité” through the “Osez l’Interdisciplinarité” initiative. CD, EM and SG were funded by ONCFS. CEFE and CROC were funded by CIL&B, MTES (ITTECOP) and FRB through the research program ERC-Lynx. CROC was funded by the European Union within the framework of the Operational Program FEDER-FSE “Lorraine et Massif des Vosges 2014–2020”, the “Commissariat à l’Aménagement du Massif des Vosges” for the FNADT (“Fonds National d’Aménagement et de Développement du Territoire”), the DREAL Grand Est (“Direction Régionale pour l’Environnement, l’Aménagement et le Logement”), the “Région Grand Est”, the “Zoo d’Amnéville”, the “Fondation d’entreprise UEM”, the “Fondation Nature & Découvertes”, the “Fondation Le Pal Nature”, and the *Chasseur d’images* magazine. This study could not have been conducted without authorizations from and agreements with municipalities, environmental managers and owners.

## Author contributions

OG wrote the paper and all co-authors commented on the manuscript. OG and SG analyzed the data. AL, CD, EG, EM and SG coordinated the study designs, the data collection and interpretation, with help from FZ for setting the experimental design in the Jura mountain.

## Data accessibility

The Eurasian lynx is an endangered species with high conservation stakes. Interactions with human activities are problematic and lead to poaching and anthropogenic pressures. Providing accurate information on lynx locations can be detrimental to the conservation status of the species. As a consequence, the original data could not be shared.

